# Transposable elements in individual genotypes of *Drosophila simulans*

**DOI:** 10.1101/781419

**Authors:** Sarah Signor

## Abstract

Transposable elements are mobile DNA sequences that are able to copy themselves within a host’s genome. Within insects they often make up a substantial proportion of the genome. While they are the subject of intense research, often times when copy number is estimated it is estimated only at the population level, or in a limited number of individuals within a population. However, an important aspect of transposable element spread is the variance between individuals in activity. Do transposable elements accumulate at different rates in different genetic backgrounds? Using two populations of *Drosophila simulans* from California and Africa I estimated transposable element copy number in individual genotypes. Some active transposable elements seem to be a property of the species, while others of the populations. I find that in addition to population level differences in transposable element load certain genotypes accumulate transposable elements at a much higher rate than others. Most likely active transposable elements are fairly rare, and were inherited only by specific genotypes that were used to create the inbred lines. Whether or not this reflects dynamics in natural populations, where transposable elements may accumulate in specific genotypes and maintain themselves in the population rather than being active at low levels population wide, is an open question.

## Introduction

Transposable elements are mobile DNA sequences that are able to move around the genome, potentially increasing their copy number in the process. They are abundant and contribute substantially to genetic variation in populations, influencing genome evolution (Biémont *et al.*, 1999; Bennetzen, 2000; Feschotte, 2008). The number and location of transposable element insertions can vary substantially between species, populations, and individuals (Vieira *et al.*, 1999; Vieira & Biémont, 2004; C Vieira, 2008; Kofler & Schlötterer, 2015; Kofler *et al.*, 2015b; Jakšić *et al.*, 2017; Kofler *et al.*, 2018). Introgression of transposable elements between species is also common, and has been reported frequently (Kofler *et al.*, 2015a; Hill & Betancourt, 2018). In addition, they make up a substantial proportion of the genome in many invertebrates and plants (Bennetzen, 2000; Tenaillon *et al.*, 2010; Kofler *et al.*, 2015b). Transposable elements are divided into three categories: long terminal repeat retrotransposons, short interspersed nuclear element retrotransposons, and transposons. The latter moves through DNA excision and repair, while the former two transpose via an RNA intermediate.

The most well studied transposable element population in insects is that of *Drosophila melanogaster*. It has been reported that in *D. melanogaster* total copy number of transposable elements may be similar between individuals, but the location of insertions can vary considerably (Barrón *et al.*, 2014). Overall copy number is thought to be maintained by transposition-selection balance, though new transposable elements may initially have high transposition rates as the host machinery evolves new defenses (Yang & Nuzhdin, 2003; Pasyukova, 2004; Slotkin & Martienssen, 2007; Johnson, 2010; Lee & Langley, 2012; Romero-Soriano & Garcia Guerreiro, 2016). Transposable elements also accumulate in heterochromatic regions, potentially because these regions are thought to be ‘transcriptionally silent’, and thus the deleterious effects of their insertion are mitigated (Dimitri, 2003). However, they can also contribute to variation in quantitative traits, differences in fitness, and changes in gene expression (Mackay 1984; Mackay 1989, Shrimpton et al. 1990, Mackay et al. 1992, Long et al. 2000). Some transposable elements have been associated with increases in fitness due to changes in gene regulation (Aminetzach *et al.* 2005; Mateo *et al.*, 2014)).

The recent explosion of population level genomic data has greatly increased our ability to estimate transposable element load in populations. Much of this work on transposable element copy number has occurred in *Drosophila*, where the existence of multiple sequenced inbred panels lend themselves to estimating copy number and insertion site frequency. However, differences in the number of transposable element insertions are thought to accumulate in inbred laboratory lines because active copies of transposable elements are rare and will be inherited only in some lines (Nuzhdin *et al.*, 1997). This is potentially mirroring what occurs in natural populations, where transposable element site heterogeneity may potentially be due to frequent transpositions in rare flies rather than low levels of transposition population-wide (Nuzhdin, 2000). Many studies of transposable element insertions estimate population-level insertion rate using Pool-seq. While Pool-seq is an effective tool for estimating population-level frequency, understanding the dynamics of transposable element spread requires the ability to estimate the variance between genotypes in transposable element copy number, in addition to population-level variation. Here I address this question by using previously sequenced lines of *D. simulans* from two populations to directly estimate transposable element copy number in individual genotypes. I estimate variance in transposable element copy number between inbred genotypes, differences between wild and inbred lines, and differences between the populations in the mean and variance of transposable element copy number.

## Methods

### Fly lines

21 African *D. simulans* isofemale lines were collected by William Ballard in 2002 from Madagascar and Peter Andolfatto in 2006 from Kenya (Jackson *et al.*, 2017). They were inbred in the lab for nine generations, during which time five were lost and DNA was collected from the initial stocks. These will be used as an estimate of ‘wild’ *Drosophila* transposable element load, compared to inbred lines. The raw reads are 90 bp paired end illumina sequencing and they were downloaded from SRA PRJEB7673 (Jackson *et al.*, 2017). The first read from each pair was used for mapping. The California lines were collected from the Zuma Organic Orchard in Los Angeles, CA on two consecutive weekends of February 2012, from a pile of rotting strawberries (Signor *et al.*, 2017; Signor & Nuzhdin, 2018; 2019). Reads were single end 100 bp and this project has been deposited at the SRA under accession SRP075682.

### Mapping and copy number estimation

Reads were mapped using bwa mem version 0.7.15 to the *D. simulans* 2.02 assembly and the 179 consensus transposable element sequences from EMBL, downloaded from Flybase.org (Li 2015). Of these 128 were used for the analysis, removing those from non-*D. melanogaster* species that did not have a presence in *D. simulans*. Bam files were sorted and index with samtools v.1.9 and optical duplicates were removed using picard MarkDuplicates (http://picard.sourceforge.net) (Li *et al.*, 2009; McKenna *et al.*, 2010). Reads with a mapping quality of below 15 were removed (this removes reads which map equally well to more than one location). Using read coverage to determine copy number has been compared to other methods and is neither permissive nor conservative (Srivastav & Kelleher, 2017). Transposable element copy number was estimated per family by comparing average coverage of the transposable sequence to average coverage of chromosome arm 2L in R, and computing population averages and individual copy number (Hill *et al.*, 2015). P-values of comparisons between means and variances were corrected for multiple testing using Bonferroni correction.

### SNPs and summary statistics

I called SNPs within the transposable elements and the genomes using GATK Haplotypecaller (McKenna *et al.*, 2010). SNPs were filtered for a minimum depth of four. SNPs were not filtered for missing calls given that not at all individuals will share insertions. Tajima’s *D* was estimated in windows of 1 kb using VCFtools, and prior to estimation indels and SNPs with more than two alleles were removed (Danecek *et al.*, 2011). The site frequency spectrum of SNPs was calculated in R to better understand insertion dynamics of transposable elements in *D. simulans*.

## Results and Discussion

### Population-level variation

Of the 128 elements examined in the population, 85 have different mean numbers of insertions between the two populations (t-test, Bonferroni corrected *p* = .05/128). Of those, only 17 are higher in the African populations, suggesting that overall the CA population has more transposable element insertion sites. Indeed, overall Californian *D. simulans* have an average of 1,797 insertions per genotype, while African *D. simulans* have 1,496. The five elements with the largest difference in copy number in Californian *D. simulans* compared to African are the *INE-1, Tc1, transib2, 1360*, and *Cr1a*. These are present on average in 37 more copies in Californian *D. simulans*. The elements more common in Africa include *baggins, X-element, R1A1-element, diver2, Rt1c, Rt1b, jockey, ninja, Osvaldo, gypsy, Bari1, gypsy8, Tirant, mariner, BS3, G5A*, and *aurora-element. G5A, gypsy8, X-element, baggins, R1A1-element, copia, G7, invader6, diver, hobo, invader3, Rt1b, Rt1c, Bari1*, and *Tirant* also have significantly different (and higher) variance compared to Californian *D. simulans* (F-test, Bonferroni corrected *p* = 0.05/128). 20 elements have different and larger variance in the Californian *D. simulans* compared to African (F-test, Bonferroni corrected *p* = 0.05/128). This includes *Stalker4, Tc3, frogger, F-element, 412, accord, G4, rover, 17.6, HMS-Beagle, gypsy6*. The most abundant transposable elements in the California population were the highly abundant *INE-1, Cr1a, G6, 1360, transib2, Tc1, baggins*, and *roo*. In the African populations this is similar, *INE-1, Cr1a, G6, baggins, R1A1-element, diver2, X-element*, and *HB*, followed by *1316* and *roo*.

*Stalker4, Stalker, Bari2, Tc3, G7*, and *Tart-C* were never present as more than a fraction of an element in any individual, and in general coverage was limited to small regions within these elements. This suggests that if these elements are present they are old and degraded. *G3* and *hopper2* are estimated as being present in ∼1 copy per individual in both populations, however closer inspection reveals that all reads are mapping to the start and end of the elements. This suggests that the only copy or copies present have internal deletions. For the *G-element* all but a small fraction of reads map to one 140 bp sequence, but they map abundantly to that particular sequence, such that the copy number is estimated as close to 6 per individual. Again, reads which map equally well to more than one location were filtered out, thus this does not represent non-specific mapping to repetitive elements. These patterns are not seen in the other transposable elements, where coverage is even across the length of the element.

The *D. melanogaster pogo* and *Helitron* elements were not present in these populations, which has been previously noted, suggesting that these transposable elements are not present in *D. simulans* (Kofler *et al.*, 2015b). A full-length version of *Quasimodo* (two copies) and *gypsy6* (one copy) were present in one individual. In other individuals reads map to very specific small regions of the transposable elements, suggesting old and degraded copies. *Stalker3* is also present in one genotype as a full-length copy. There is no mapping in the African populations and very little in Californian populations, suggesting that old or degraded copies are not present in the rest of the genotypes sampled.

### Site frequency spectrum

I examined the site frequency spectrum of each transposable element in African and Californian *D. simulans* (Table 2). In some cases there are no polymorphisms (*Dmau\mariner*) or very few (*Dmel\p-element*) therefore this is uninformative. Furthermore, overall the Californian population has more intermediate frequency polymorphisms (measured using Tajima’s *D*; Signor et al. 2017) compared to the African population (Supplementary Figure 1), which may be expected to affect the site frequency spectrum overall. There are different amounts of information in these estimates, thus they must be interpreted along with Table 1 – for example *Quasimodo* is really on present in two full length copies in a single individual, thus this estimation of the site frequency spectrum is not informative with regards to the spread of *Quasimodo* in the population.

**Table 1:**
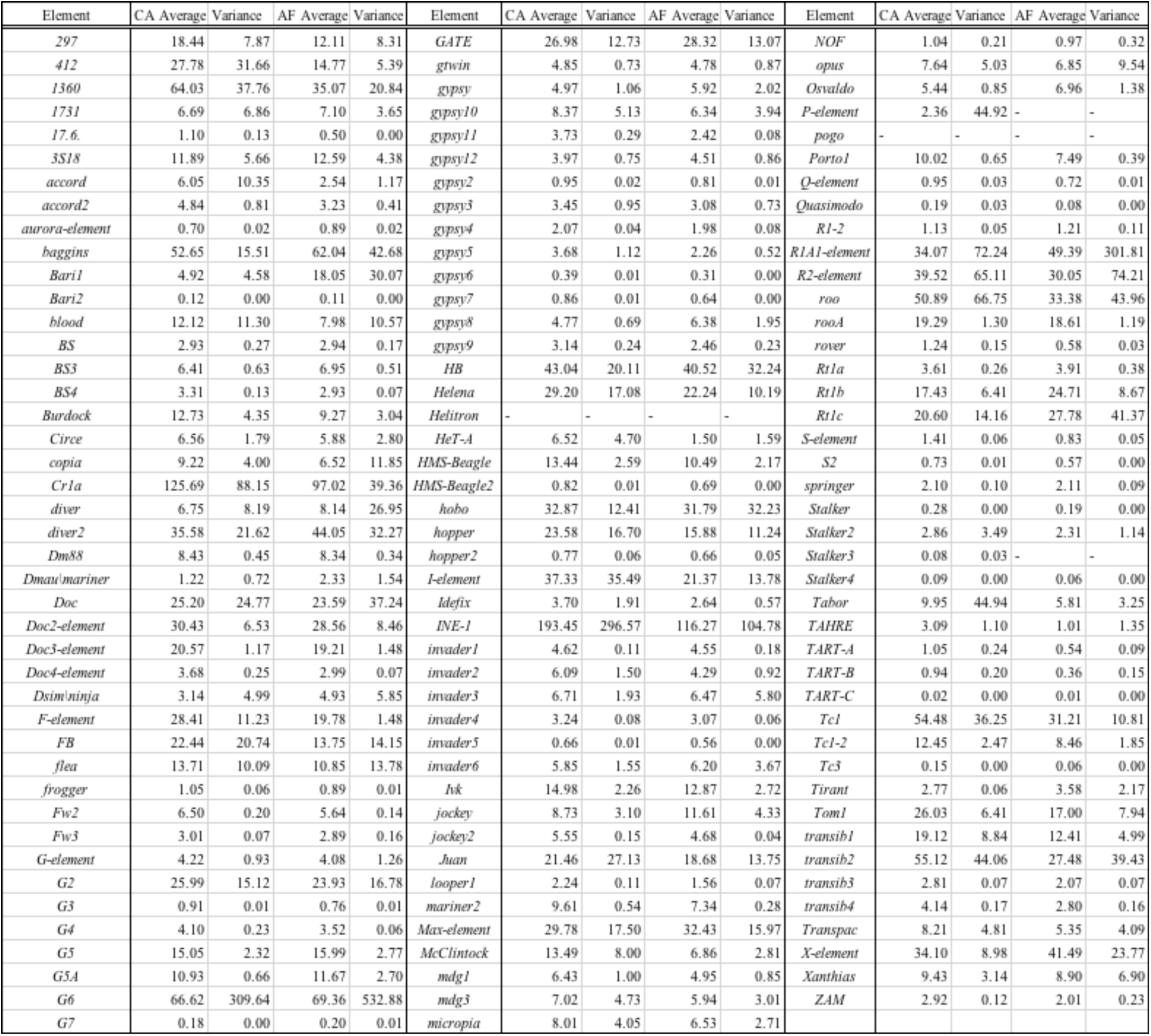
Summary statistics for copy number of transposable elements from the Californian and African *D. simulans*.

**Table 2:**
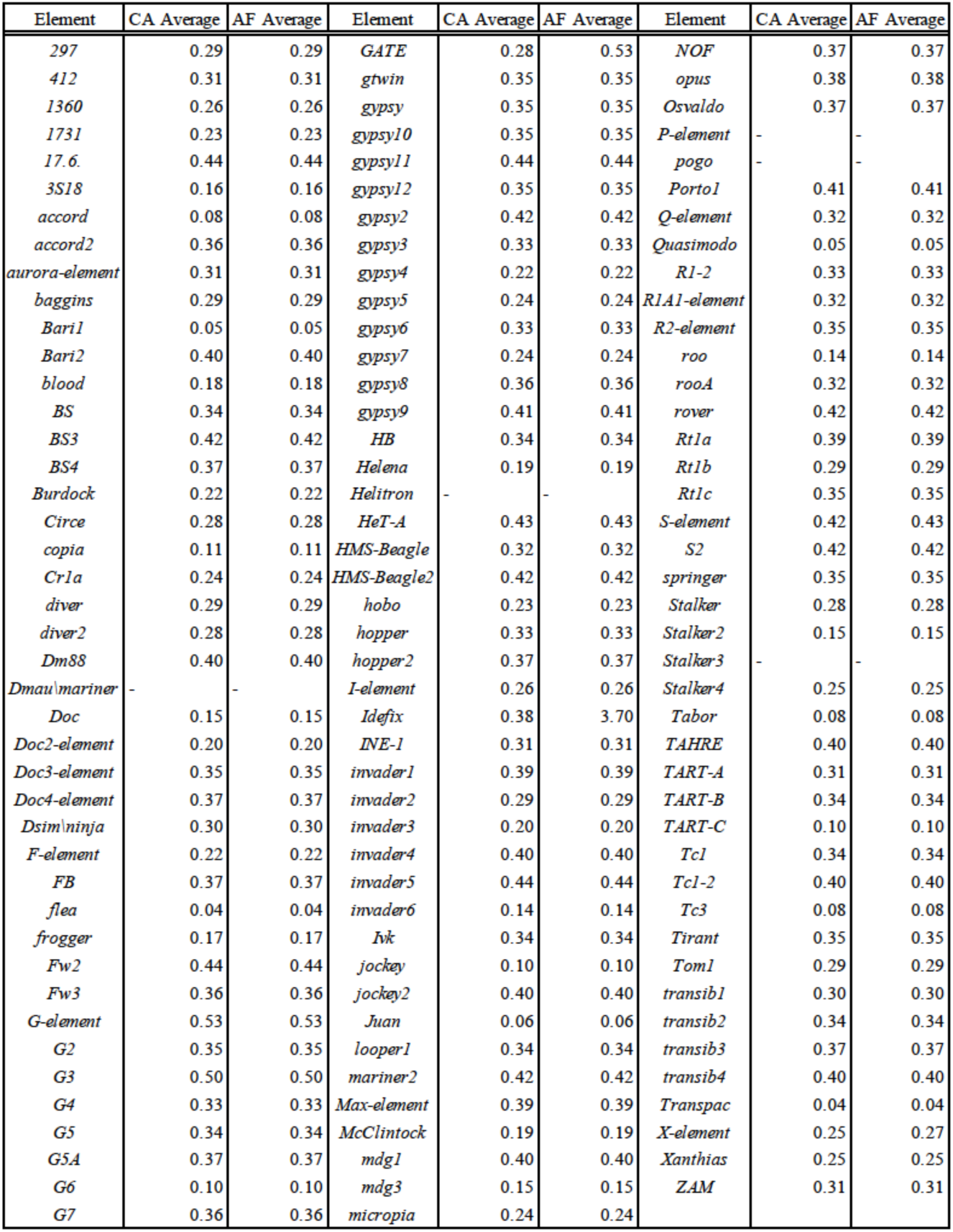
Average site frequency spectrum for each element, excluding sites that are fixed relative to *D. melanogaster. Dmau\mariner* and the *p-element* are present in at least one population, but have no polymorphic SNPs. Other elements without an estimated site frequency spectrum are not present in the population.

**Figure 1:**
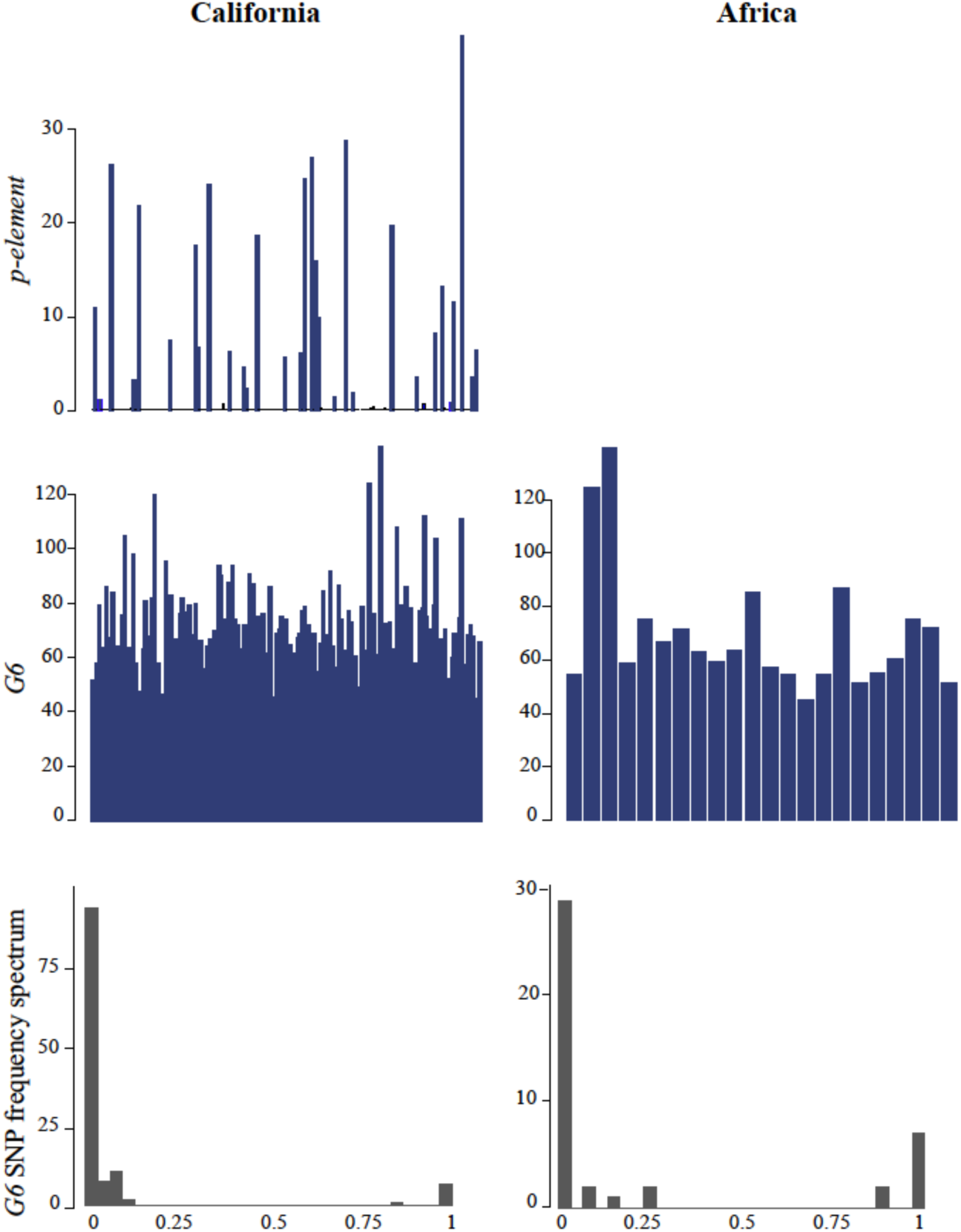
Estimated copy number for the *p-element* and *G6* in Californian and African *D. simulans*. Each bar represents an individual from the population. As expected, the *p-element* was not found in the African population sampled in the early 2000’s, but by 2012 when the Californian *D. simulans* was sampled it had invaded. *G6* has a high copy number in both populations of *D. simulans*, which wasn’t recorded in previous studies on African *D. simulans*. The site frequency spectrum in the last row also suggests recent spread of the *G6* element in *D. simulans*, as there are primarily low frequency SNPs. Note that while fixed SNPs are included in this graph to illustrate divergence from D. melanogaster they are not included in the estimation of average site frequency spectrum shown in Table 2.

**Figure 2:**
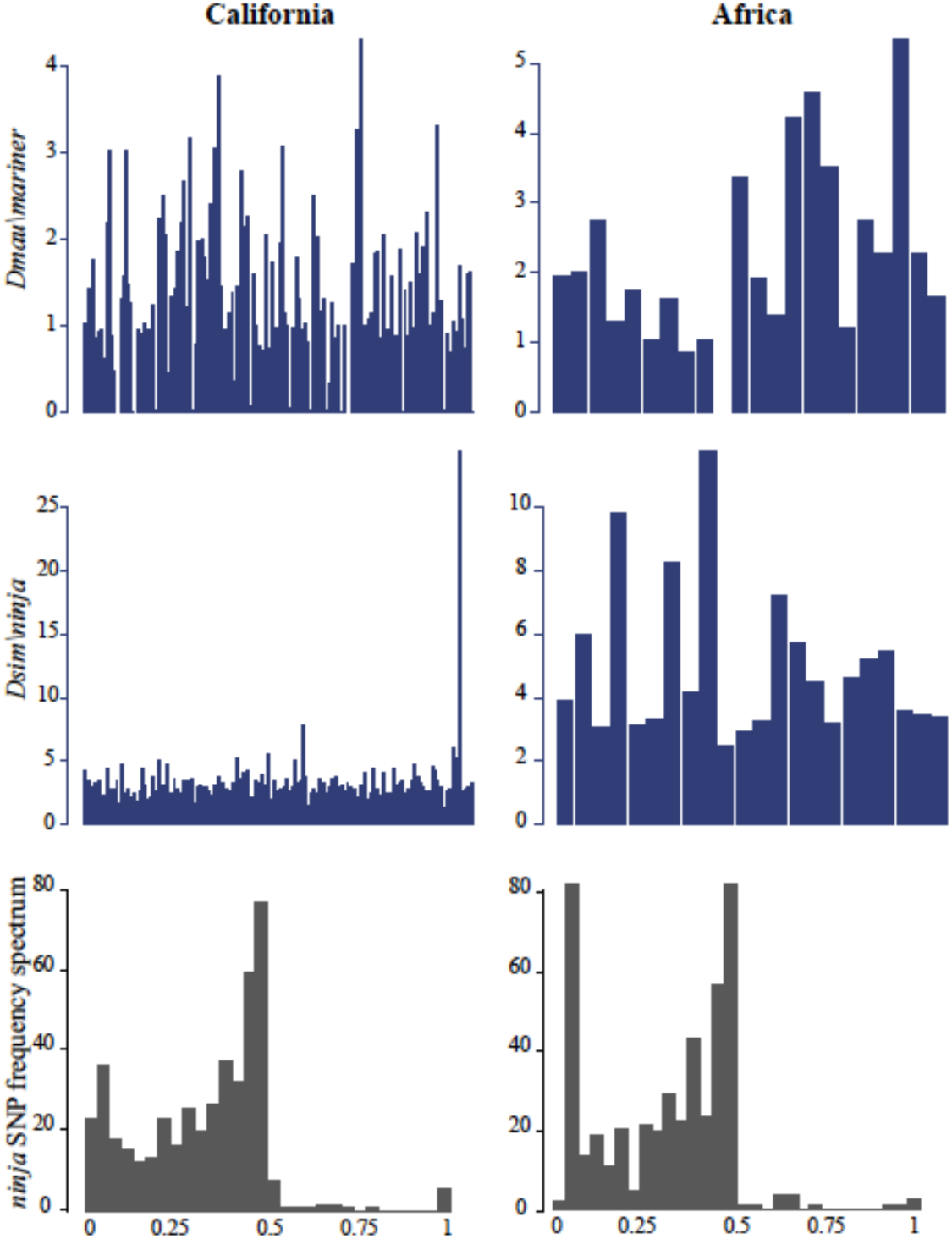
Estimated copy number for the two non-*D. melanogaster* transposable elements included here, *Dsim\ninja* and *Dmau\mariner*. Variance between individual genomes is large for transposable elements, for example in California *Dsim\ninja* varies from 1-29, and in Africa from 2-12. The individual with 29 copies of *Dsim\ninja* has 10 fixed SNPs within the element and 27 polymorphisms compared to other population members (for example, 239 polymorphisms and 5 fixed SNPs), suggesting that it has been recently active within that genome. The site frequency spectrum of *D. sim\ninja* is broad, suggesting that outside of the genome with an active copy of *D.sim\ninja* this element has been diverging within this species for some time. In contrast, no SNPs were called in *Dmau\mariner*, suggesting recent colonization in *D. simulans*.

Elements with site frequency spectrum heavily biased towards low frequency SNPs in Californian *D. simulans* include *G6, flea*, and *Juan* (Figure 1, Table 2). In African *D. simulans* this includes *Tabor, Transpac, flea, Juan, Bari1, G6*, and *accord* (Figure 1, Table 2). Thus in both populations *G6, flea, Bari1*, and *Juan* likely have recent activity. This is consistent with other work on *Juan*, which suggests it is actively transposing in the species (Kofler *et al.*, 2015b). The larger number of transposable elements with low frequency SNPs in African populations may be due to the overall difference in the site frequency spectrum between populations (Supplementary Figure 1; Signor *et al.* 2017).

### The p-element

The *p-element* recently invaded *D. simulans* from *D. melanogaster* as described in (Kofler *et al.*, 2015a), however Pool-seq cannot tie *p-element* insertions to specific individuals, only determine the average number of insertions. What was reported previously was 0.4 insertions in Florida populations, and 29 in South Africa (Kofler *et al.*, 2015a). What we see in the California population is an average of 2 insertions, however that is because the majority of individuals do not have any insertions (137 individuals have less than .3 estimated copies, Figure 1). The remaining individuals have between .5 and 39 copies. We recovered the single reported SNP in the *p-element*, a G to A transition at position 2040. It is interesting that it is not invading genotypes in the population at the same rate, but rather reaching high copy number in some genotypes and not others (Nuzhdin 2000). However, given the observed ‘outliers’ (see next section), one genotype for a given transposable element with high copy number, this may be reflective of how some transposable elements invade populations. It is also possible that *p-elements* are just proliferating in strains that contained an active copy prior to collection (Nuzhdin & Mackay 1997). This was observed previously in lab strains of *D. melanogaster*, though contamination and introgression may also have played a role (Rahman *et al.* 2015).

### Transposable elements in individual genomes

Some transposable elements have considerably higher copy number in particular genotypes compared to the population average. For example, in one genotype *Dsim\ninja* is present in 29 copies, compared to the population mean of three. *Dsim\ninja* has 10 fixed differences and 27 polymorphisms in this strain from the California population, and the population average is 7.5 fixed differences and 264 polymorphisms (Supplementary Figure 4 & 5). This suggests that *Dsim\ninja* was recently active in this genome. This is true of several transposable elements which have outliers in the population. *Stalker2* has an outlier with 17 fixed SNPs and 8 polymorphisms, compared to a population average of 14 fixed SNPs and 43 polymorphisms. The *gypsy10* element has a single outlier individual with 203 fixed differences compared to *D. melanogaster*, and 77 polymorphisms. The population mean is 158 fixed differences and 222 polymorphisms. Other transposable elements with large outliers in the California population include *opus, blood, GATE, diver, Tabor, INE-1, diver2, idefix, 1731, 412*, and *297*.

Sampling of the African populations was much more limited thus less genotype-specific variation is sampled, and indeed only two transposable elements had large outliers, both in the same individual from Madagascar: *copia*, and *diver*. This individual had 11 fixed differences in *copia*, and 20 polymorphisms, while the population average is 10 fixed differences and 78 polymorphisms (and 20 copies compared to 4-11 for the rest of the population). For *diver* this individual had 20 fixed differences and 84 polymorphisms, compared to a population average of 12 fixed differences and 215 polymorphisms (and 30 copies compared to 4-10 for the rest of the population). The outlier individual is one that was inbred in the lab. In general being inbred in the lab is not affecting overall transposable element copy number however, as comparing between lines that were sequenced directly upon collection and those that there inbred, there is no significant difference between the mean number of transposable elements for any transposable element family. All of this is to suggest that transposable element load varies significantly between genotypes within a population, though which transposable elements are able to ‘wake up’ and multiply may be shared between genotypes in different populations (*diver*). That is to say that some recent transposable element activity is shared across the species (*G6*), populations (*p-element*) and some is specific to genotypes within a population. Those that ‘wake up’ in individual lines appears to be due to sampling of individuals that are permissive or contain active transposable elements, rather than an overall increase in transposable element activity in inbred lines.

### Comparison to other studies

Previously *HMS Beagle, blood, flea, opus, Stalker*, and *F-element* were described as being present in 0-1 copies in most populations from a large survey, with some populations having a significant number of insertions (Biémont *et al.*, 1999). In our populations all of these elements with the exception of *Stalker* were present in on average more than six insertions. Of these only *flea* has more low frequency polymorphisms suggesting that it has spread within the genome recently. Vieira *et al.*, 1999 detected *17.6* at one insertion site in *D. simulans*, and here it was present on average in 1 (CA) to 0.5 (AF) copies in these populations, which is more frequent than recent papers such as Kofler *et al.*, 2015b which detected four copies in the population of ∼800 which was sampled. Vieira *et al.*, 1998 estimated 0-30 copies of *412* and 20-100 copies of *roo* per genome including heterochromatin. We observed 5-42 copies of *412* in both populations, with a single outlier at 73. For *roo* we observed 20-75 copies in both populations. Thus our estimates are consistent with previous estimates for these elements that include heterochromatin. It has also been suggested that *412* is recently active (Biémont *et al.*, 1999).

*Tirant* has previously been reported as having higher copy number in African *D. simulans*, potentially due to a recent mobilization of the element (Fablet *et al.*, 2006). We find that pattern here, including a higher variance in the African populations where copy number ranges from 2 to 6.68, compared to 2-3.8 in California (Fablet *et al.*, 2006). Viera et al. 1999 sampled diverse *D. simulans* populations and found ∼1 *Tirant* element per genome, though this was limited to euchromatic regions and older *Tirant* insertions may potentially have accumulated in heterochromatic regions. The *Dmau\mariner* element has a higher copy number in Africa than in the Californian *D. simulans*, from 0-5 with an average of 2.33, compared to 0-3 with an average of 1.22. *Dmau\mariner* also contains no polymorphisms, which is consistent with previous work which found sequence identify between *mariner* copies of ∼99% (Capy et al. 1990; Capy et al. 1992). This is also consistent with a recent spread of *Dmau\mariner* in *D. simulans* (Capy et al. 1990; Capy et al. 1992). For *mariner2* we find copy number to be relatively stable in both populations (9.6 in California, 7.3 in Africa), while in Kofler et al. 2015 *mariner2* is only reported in six copies in the entire population of ∼800, though again this is limited to euchromatic regions.

The *G6* element has a large difference from previously reported values, with an average of 66 insertions in Californian *D. simulans* and 69 in African. However, only 37 insertions where reported total for a previously estimated population of ∼800 *D. simulans* isofemale lines (Kofler *et al.*, 2015b). Only 18% of reads in Kofler et al. (2015) map to euchromatin, but this is not a large enough fraction to explain the difference. In addition, in the populations reported here the *G6* element has primarily low frequency polymorphisms (Table 2), suggesting that this is a recent expansion of copy number. The population examined in Kofler et al. (2015) is from South African in 2013, a later collection date than either population here, though it is possible that the *G6* activation is geographically limited to more northern populations. Overall our estimates are higher than the work of Kofler et al. 2015, which only estimates more than one insertion per line for four transposable elements (*1360, hobo, roo* and *Tc-2*). This is conservative compared to other estimates of euchromatic insertions, for example Vieira et al. 1999 use *in situ* hybridization to estimate copy number and found more than one insertion on average for nineteen elements (of 36 measured, *412, blood, burdock, copia, coral, flea, gypsy, HMS beagle, mdg3, opus, roo, Tirant, dor, F-element, Helena, I-element, jockey, Bari1*, and *hobo*). There is other evidence this work may be conservative, for example Kaminker *et al.*, 2002 estimated 146 *roo* insertions in the euchromatin of the *D. melanogaster* reference genome, while Kofler *et al.*, 2015b estimates 1966 total for a population of 554 (approximately 3.6 per individual). This may also be due to inherent variability between populations.

## Conclusions

The general picture that one draws from this summary is that non-African *D. simulans* have a higher insertion number than African populations. *D. simulans* is currently being invaded by transposable elements, and this spread is likely occurring concordant with the world-wide colonization of *D. simulans*, as has been posited by previous studies (Lachaise et al., 1988; Vieira et al., 1999; Biémont et al., 2003). However, African populations have their own transposable element dynamics, with some transposable elements seeming to share activity between populations (*G6*) and others being more active in African *D. simulans* (*baggins, Bari1*, etc.).

Transposable element load is an attribute of species, populations, and individual genotypes. In inbred laboratory lines active transposable element copies may be inherited by some lines and not others, and active transposable elements can accumulate over time (Nuzhdin *et al.*, 1997). This can cause differences over time in the number of insertions within a line and large differences between lines in transposable element copy number (Nuzhdin *et al.*, 1997). This may also be reflective of natural patterns in which transposable elements proliferate in particular genotypes rather than at low levels in the population as a whole (Nuzhdin, 2000). Overall, looking at variance between individuals is an important part of understanding the ways in which transposable elements maintain themselves in populations.

## Acknowledgements

I would like to thank C. and S. Emery for helpful commentary on the manuscript. I am also thankful to J. Butler and T. Robert for help in the laboratory

The author declares that she has no conflicting interests.

## References

Aminetzach YT, Macpherson JM, Petrov DA. 2005. Pesticide resistance via transposition-mediated adaptive gene truncation in *Drosophila*. Science 309:764–7.

Barrón, M.G., Fiston-Lavier, A.-S., Petrov, D.A. & González, J. 2014. Population genomics of transposable elements in *Drosophila*. Annu. Rev. Genet. 48:561–581.

Bennetzen, J.L. 2000. Transposable element contributions to plant gene and genome evolution. Plant Mol. Biol. 42:251–269.

Biémont, C., Vieira, C., Borie, N. & Lepetit, D. 1999. Transposable elements and genome evolution: the case of *Drosophila simulans*. Genetica 107:113–120.

Biémont C, Nardon C, Deceliere G, Lepetit D, Lœvenbruck C, Vieira C. 2003. Worldwide distribution of transposable element copy number in natural populations of *Drosophila simulans*. Evolution 57:159–67.

Capy P, Chakrani F, Lemeunier F, Hartl DL, David JR. 1990. Active *mariner* transposable elements are widespread in natural populations of *Drosophila simulans*. Proc R Soc B 242:57–60.

Capy P., Koga A., David J.R., Hartl D.L. 1992. Sequence analysis of active *mariner* elements in natural populations of *Drosophila simulans*. Genetics 130:499–506.

Danecek, P., Auton, A., Abecasis, G., Albers, C.A., Banks, E., DePristo, M.A., et al. 2011. The variant call format and VCFtools. Bioinformatics 27:2156–2158.

Dimitri, P. 2003. Colonization of heterochromatic genes by transposable elements in *Drosophila*. Mol. Biol. Evol. 20:503–512.

Fablet, M., McDonald, J.F., Biémont, C. & Vieira, C. 2006. Ongoing loss of the *tirant* transposable element in natural populations of *Drosophila simulans*. Gene 375:54–62.

Feschotte, C. 2008. Transposable elements and the evolution of regulatory networks. Nat. Rev. Genet. 9:397–405.

Hill, T. & Betancourt, A.J. 2018. Extensive exchange of transposable elements in the *Drosophila pseudoobscura* group. 1–14. Mobile DNA.

Hill, T., Schlötterer, C. & Betancourt, A. 2015. Hybrid dysgenesis in *Drosophila simulans* due to a rapid global invasion of the P-element. PLoS 12:e1005920.

Jackson, B.C., Campos, J.L., Haddrill, P.R., Charlesworth, B. & Zeng, K. 2017. Variation in the intensity of selection on codon bias over time causes contrasting patterns of base composition evolution in *Drosophila*. Genome Biol. Evol. evw291–22.

Jakšić, A.M., Kofler, R. & Schlötterer, C. 2017. Regulation of transposable elements: interplay between TE-encoded regulatory sequences and host-specific trans-acting factors in *Drosophila melanogaster*. Mol. Ecol. 26:5149–5159.

Johnson, N.A. 2010. Hybrid incompatibility genes: remnants of a genomic battlefield? Trends Genet. 26:317–325.

Kaminker, J.S., Bergman, C.M., Kronmiller, B., Carlson, J., Svirskas, R., Patel, S., et al. 2002. The transposable elements of the *Drosophila melanogaster* euchromatin: a genomics perspective. Genome Biology 3:research0084.1.

Kofler, R. & Schlötterer, C. 2015. Low levels of transposable element activity in *Drosophila mauritiana*: causes and consequences. bioRxiv.

Kofler, R., Hill, T., Nolte, V., Betancourt, A.J. & Schlötterer, C. 2015a. The recent invasion of natural *Drosophila simulans* populations by the *P-element*. Proc. Nat. Acad. Sci. USA 112:6659–6663.

Kofler, R., Nolte, V. & Schlötterer, C. 2015b. Tempo and mode of transposable element activity in *Drosophila*. PLoS Genet. 11:e1005406.

Kofler, R., Senti, K.-A., Nolte, V., Tobler, R. & Schlötterer, C. 2018. Molecular dissection of a natural transposable element invasion. Genome Res. 28:824–835.

Lachaise, D., M. Cariou, J. R. David, F. Lemeunier, L. Tsacas, and M. Ashburner. 1988. Historical biogeography of the *Drosophila melanogaster* species subgroup. Evol. Biol. 22:159–227.

Lee, Y.C.G. & Langley, C.H. 2012. Long-term and short-term evolutionary impacts of transposable elements on *Drosophila*. Genetics 192:1411–1432.

Li, H., Handsaker, B., Wysoker, A., Fennell, T., Ruan, J., Homer, N., et al. 2009. The Sequence Alignment/Map format and SAMtools. Bioinformatics 25:2078–2079.

Li, H. 2015. Aligning sequence reads, clone sequences and assemble contigs with BWA-MEM arXiv 1–3.

Long, A.D., Lyman, R.F., Morgan, A.H., Langley, C.H., Mackay, T.F. 2000. Both naturally occurring insertions of transposable elements and intermediate frequency polymorphisms at the *achaete-scute* complex are associated with variation in bristle number in *Drosophila melanogaster*. Genetics 154:1255–69.

Mackay, T.F.C. 1984. Jumping genes meet abdominal bristles: hybrid dysgenesis-induced quantitative variation in *Drosophila melanogaster*. Genet. Res. 44:231–237.

Mackay, T.F.C. 1989. Transposable elements and fitness in *Drosophila melanogaster*. Genome 31:284–295.

Mackay, T.F., Lyman, R.F., Jackson, M.S., 1992. Effects of P element insertions on quantitative traits in *Drosophila melanogaster*. Genetics 130:315–332.

Picot, S., Wallau, G.L., Loreto, E.L.S., Heredia, F.O., Hua-Van A., Capy P. 2008. The *mariner* transposable element in natural populations of *Drosophila simulans*. Heredity 101:53–9.

Rahman R., Chirn, G.-W., Kanodia, A., Sytnikova, Y.A., Brembs, B., Bergman, C.M., et al. 2015. Unique transposon landscapes are pervasive across *Drosophila melanogaster*. Nucleic Acids Res. 43:10655–72.

Mateo, L., Ullastres, A. & González, J. 2014. A Transposable element insertion confers xenobiotic resistance in *Drosophila*. PLoS Genet. 10:e1004560.

McKenna, A., Hanna, M., Banks, E., Sivachenko, A., Cibulskis, K., Kernytsky, A., et al. 2010. The Genome Analysis Toolkit: a MapReduce framework for analyzing next-generation DNA sequencing data. Genome Res. 20:1297–1303.

Nuzhdin, S.V. 2000. Sure facts, speculations, and open questions about the evolution of transposable element copy number. In: Transposable Elements and Genome Evolution (J. F. McDonald, ed), pp. 129–137. Springer Netherlands, Dordrecht.

Nuzhdin, S.V., Pasyukova, E.G. & Mackay, T.F. 1997. Accumulation of transposable elements in laboratory lines of *Drosophila melanogaster*. Genetica 100:167–175.

Pasyukova, E.G. 2004. Accumulation of transposable elements in the genome of *Drosophila melanogaster* is associated with a decrease in fitness. Journal of Heredity 95:284–290.

Romero-Soriano, V. & Garcia Guerreiro, M.P. 2016. Expression of the retrotransposon *Helena* reveals a complex pattern of TE deregulation in *Drosophila* hybrids. PLoS ONE 11:e0147903.

Signor, S. & Nuzhdin, S. 2018. Dynamic changes in gene expression and alternative splicing mediate the response to acute alcohol exposure in *Drosophila melanogaster*. Heredity 121: 342-360.

Signor, S.A. & Nuzhdin, S.V. 2019. Evolution of phenotypic plasticity in response to ethanol between sister species with different ecological histories (*Drosophila melanogaster and D. simulans*). bioRxiv.

Signor, S.A., New, F.N. & Nuzhdin, S. 2017. A large panel of *Drosophila simulans* reveals an abundance of common variants. Genome Biol. Evol. 10:189–206.

Shrimpton, A.E., Mackay, T.F., Brown, A.J. 1990. Transposable element-induced response to artificial selection in *Drosophila melanogaster*: molecular analysis of selection lines. Genetics 125:803–11.

Slotkin, R.K. & Martienssen, R. 2007. Transposable elements and the epigenetic regulation of the genome. Nat. Rev. Genet. 8:272–285.

Srivastav, S.P. & Kelleher, E.S. 2017. Paternal induction of hybrid dysgenesis in *Drosophila melanogaster* is weakly correlated with both *P-element* and *hobo* element dosage. G3 7:1487–1497.

Tenaillon, M.I., Hollister, J.D. & Gaut, B.S. 2010. A triptych of the evolution of plant transposable elements. Trends Plant Sci. 15:471–478.

Vieira, C.B. 2008. Selection against transposable elements in *D. simulans* and *D. melanogaster*. Genet. Res. 68:9–115

Vieira, C. & Biémont, C. 2004. Transposable element dynamics in two sibling species: *Drosophila melanogaster* and *Drosophila simulans*. Genetica 120:115–123.

Vieira, C., Aubry, P., Lepetit, D. & Biémont, C. 1998. A temperature cline in copy number for *412* but not *roo*/B104 retrotransposons in populations of *Drosophila simulans*. Proc. R. Soc. B 265:1161–1165.

Vieira, C., Lepetit, D., Dumont, S. & Biémont, C. 1999. Wake up of transposable elements following *Drosophila simulans* worldwide colonization. Mol. Biol. Evol. 16:1251–1255.

Yang, H.-P. & Nuzhdin, S.V. 2003. Fitness costs of *Doc* expression are insufficient to stabilize its copy number in *Drosophila melanogaster*. Mol. Biol. Evol. 20:800–804.

