## Supplemental Files for "Transposable elements in individual genotypes of *Drosophila simulans*" (from another version of this preprint: version 1, 2019-09-25)

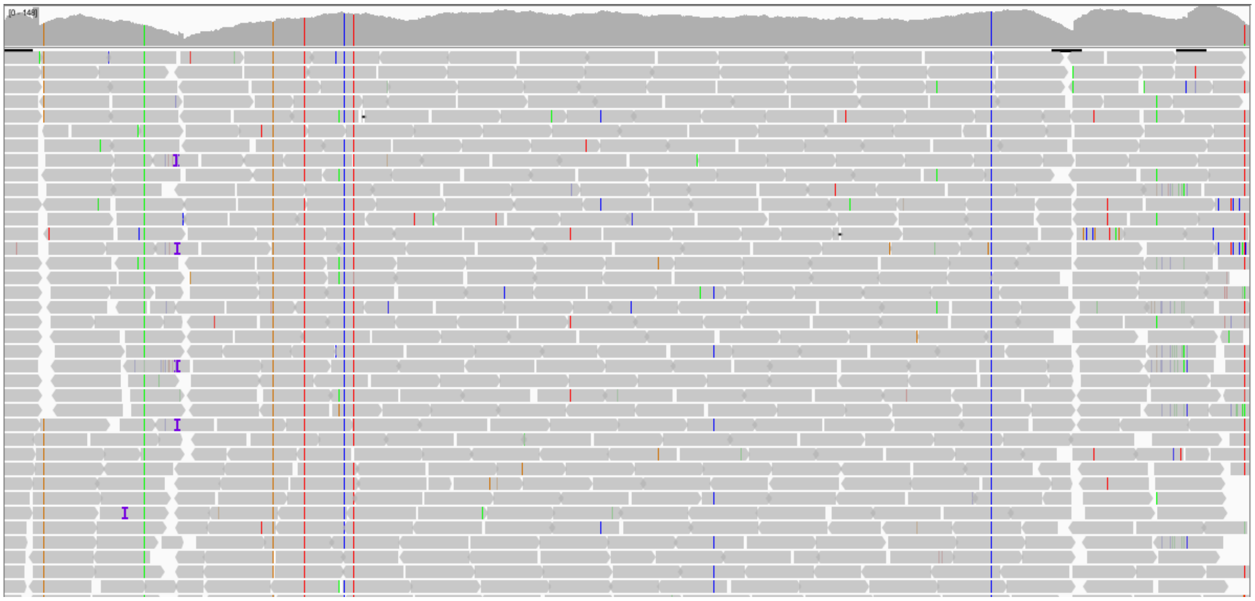

Supplementary Figure 1: Representative mapping to the *G6* element in Californian *D. simulans*. There is little variation and some low frequency SNPs, suggesting a recent spread in both the African and Californian populations.

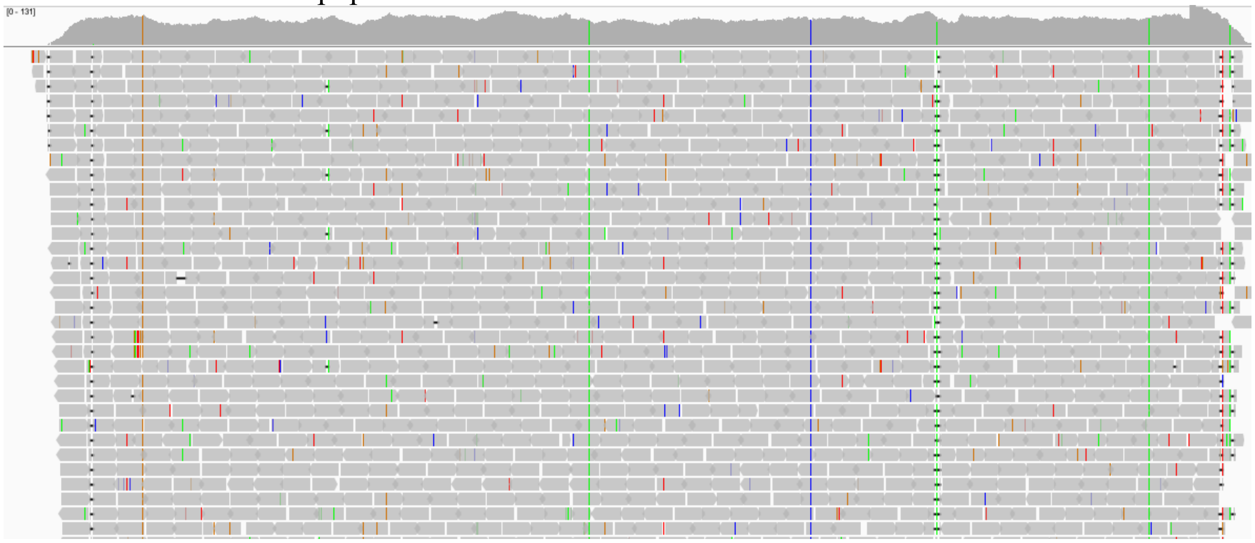

Supplementary Figure 2: Representative mapping of *flea* in the Californian population, with few SNPs and many low frequency polymorphisms, suggesting recent spread.

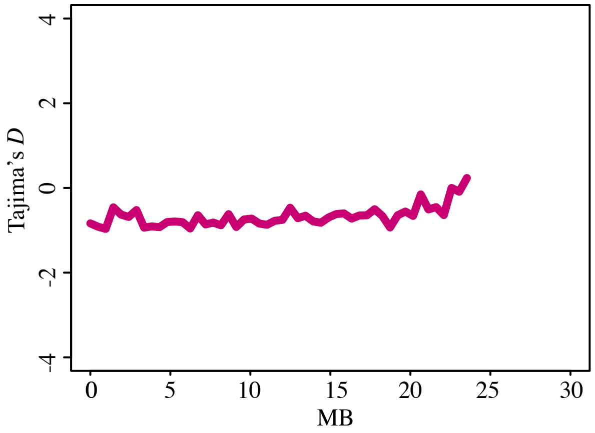


Supplementary Figure 3: Representative Tajima's *D* from 2L of the African population of *D. simulans*.

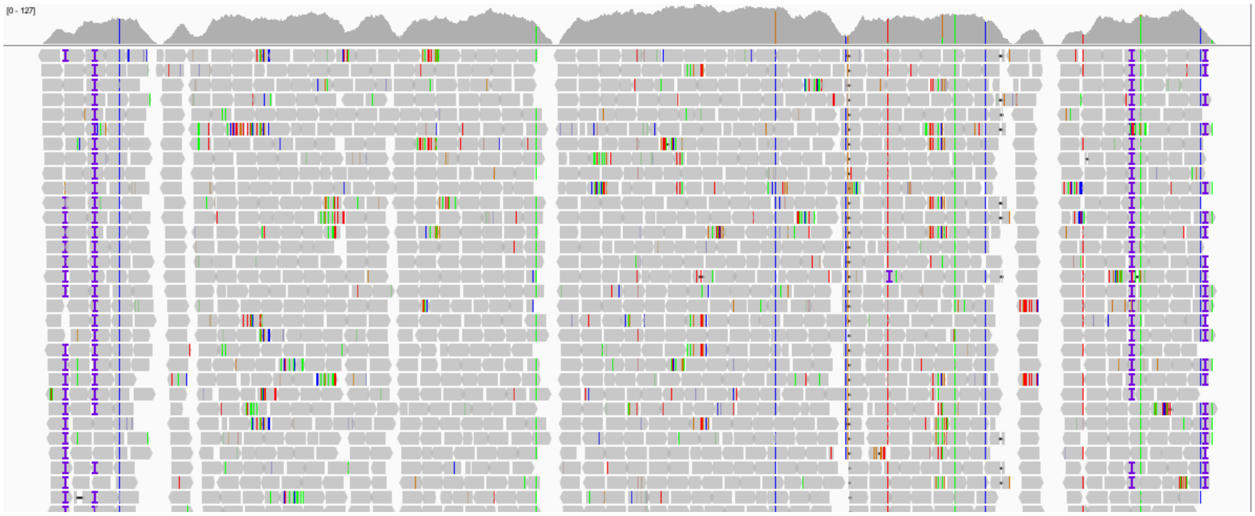

Supplementary Figure 4: Mapping of reads from one genotype with a large number of *Dsim\ninja* copies compared to the rest of the population (29 copies compared to a mean of 3). This suggest recent activity of the element in this genome.

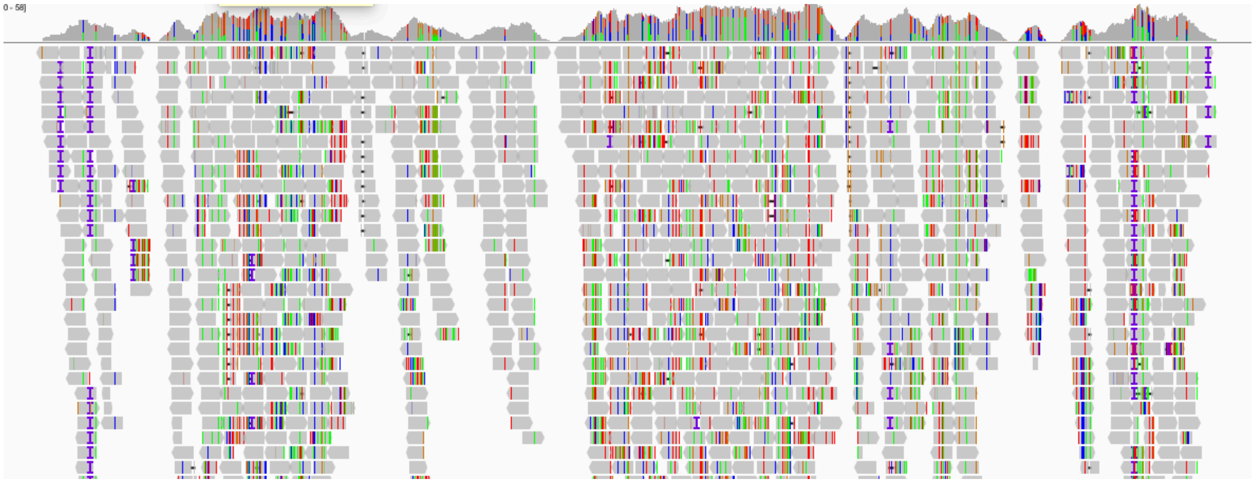

Supplementary Figure 5: For comparison, this is mapping of reads from a single genotype to *Dsim\ninja*, in an individual with copy number representative of the population mean (3).
